# A humanized ossicle platform for real-time tracking and preclinical assessment of CAR-based immunotherapies in acute myeloid leukemia

**DOI:** 10.64898/2026.06.22.733693

**Authors:** S Donsante, M Algeri, M Biondi, V Zambelli, C Guzzetti, E Grassenis, G Alberti, E Rezoagli, S Tettamanti, A Biondi, M Riminucci, A Pievani, M Serafini

## Abstract

Preclinical evaluation of chimeric antigen receptor (CAR)-T therapies for acute myeloid leukemia (AML) is limited by the lack of models that faithfully recapitulate the human bone marrow (BM) niche. Here, we implemented a humanized ossicle-based AML model that enables simultaneous engraftment of leukemic blasts and longitudinal assessment of responses to CAR-based immunotherapies. Intravenous or intra-ossicle injection of AML blasts produced robust, reproducible disease mimicking features of human AML within its microenvironment. To monitor tumor burden and immune effector cells in real-time, we developed a dual bioluminescence system using distinct luciferases in AML and CAR-T cells. This approach allowed non-invasive longitudinal tracking of CAR-T cell localization, expansion, persistence, and leukemic clearance within the ossicle. Overall, our model provides a powerful platform to study CAR-T cell behavior within a human BM niche and, for the first time, allows simultaneous longitudinal visualization of leukemic burden and CAR-T cell dynamics in a physiologically relevant ossicle-based AML model.

**Teaser:** Humanized ossicles combined with dual BLI enable tracking of AML progression and CAR-T cell dynamics in a human stromal niche.

## Introduction

Chimeric antigen receptor (CAR)-based cell therapy has revolutionized the treatment landscape of B-cell malignancies but remains largely ineffective in acute myeloid leukemia (AML) (*1*). Therapeutic failure in AML is driven by multiple factors, including heterogeneous target antigen expression, shared antigen expression between leukemic and healthy hematopoietic cells, and the complex challenges related to leukemic bone marrow (BM) microenvironment (*2*, *3*). Despite advances in target antigen selection and CAR design, clinical translation is still limited by the lack of preclinical models that accurately recapitulate human AML in its native BM niche (*4*).

Current preclinical evaluation of CAR-based immunotherapies for AML mainly depends on xenotransplantation models, which present important limitations that compromise their translational predictivity. The most critical issue is the suboptimal cross-reactivity between human hematopoietic cells and the murine microenvironment (*5*). Species-specific discrepancies in stromal cellular composition, cell-adhesion molecules, and cytokine signalling often mask the contribution of the niche to AML engraftment, progression, and therapy resistance (*6–8*).

These limitations are particularly critical for the evaluation of next-generation CAR-based therapies, which are designed not only to target tumor antigens but also to actively engage and reshape the tumor microenvironment (*9*, *10*). Such approaches, including armored CARs, cytokine-engineered constructs, and chemokine receptor–redirected cells, rely heavily on intact human stromal cues and trafficking signals that are poorly reproduced in conventional xenograft models, underscoring the critical need for more physiologically relevant humanized systems (*11*, *12*).

A promising strategy that we and others developed in recent years is the generation of ectopic humanized ossicles through the subcutaneous implantation of scaffold-free chondrogenic pellets derived from primary human BM mesenchymal stromal cells (MSCs) into immunocompromised mice. This system recreates the three-dimensional architecture, cellular heterogeneity and molecular profile of the native BM microenvironment (*13*, *14*). The resulting ossicles are composed of bony cortex, bone trabeculae and a functional marrow cavity, thus representing a faithful model of the human stromal niche that offers a significant advance over conventional xenografts. Particularly, by providing a species-matched niche, humanized ossicles enable the engraftment of human hematopoietic cells, including patient-derived AML samples that poorly engraft in mouse femurs (*15*, *16*). Despite these advantages, standard ossicle models derived from conventional cartilaginous pellets have significant practical limitations, as their small size limits their utility for comprehensive preclinical studies.

Another critical limitation of current AML models is the difficulty to longitudinally monitor both leukemic burden and CAR-engineered cell dynamics simultaneously in the same animal. While antitumor activity remains the primary endpoint in preclinical CAR-based immunotherapy evaluations, analysis of *in vivo* trafficking, homing, expansion, and long-term persistence of infused cells is equally critical as these processes, which are profoundly influenced by the leukemic microenvironment, all affect the therapeutic efficacy (*17*). Orthogonal bioluminescence imaging (BLI) systems represent high-sensitivity solutions for tracking infused cells *in vivo* (*18*), with dual-luciferase approaches enabling the simultaneous assessment of CAR-engineered cell distribution and leukemic progression within the same animal (*19–22*). This dual-imaging system offers real-time, high-resolution insights into the kinetics of therapeutic response, enabling direct visualization of effector cell localization, expansion kinetics, persistence patterns, and their relationship to leukemic clearance.

In this study, we address these key limitations in AML preclinical models through three main advances. First, we demonstrate for the first time that ossicles can be used as an experimental platform to combine AML modeling with the evaluation of CAR-based immunotherapy within a humanized BM niche. Second, we optimize the ossicle generation protocol through basement membrane extract (BME) supplementation, markedly increasing ossicle size and expanding the human BM niche while preserving its structural and biological integrity. This enhancement improves the overall efficiency of the model and enables controlled local delivery of AML cells. Third, we develop and validate a dual-luciferase imaging system that allows real-time monitoring of both anti-CD33 CAR-engineered cytokine-induced killer (CIK) cell and AML progression within the ossicle model, providing a proof-of-concept of its utility for longitudinal evaluation of CAR-based immunotherapies.

By combining optimized BME-ossicles with dual BLI, we established an advanced preclinical platform for the evaluation of anti-AML CAR-based immunotherapies and the functional assessment of next-generation CAR-engineered cells.

## Results

### Humanized ossicles support AML engraftment and enable evaluation of CAR-based immunotherapy

First, we investigated the capacity of our humanized ossicles to support the engraftment of AML cells within the reconstructed human BM microenvironment. Six weeks after chondrogenic pellet implantation, corresponding to the onset of marrow cavity formation, NSG mice were intravenously injected with AML cell lines (KG-1 and HL-60) or patient-derived xenograft (PDX) AML cells (**Figure 1A**). Histological and immunofluorescence (hCD33^+^) analysis of ossicles explanted after 2 and 4 weeks respectively revealed a well-organized marrow cavity heavily infiltrated by myeloid blasts, confirming successful AML engraftment within the ossicles (**Figure 1B**). Flow cytometric quantification demonstrated that myeloblast engraftment levels in the ossicle were comparable to, or in some instances exceeded, those observed in native murine BM (**Figure 1C**).

**Figure 1.**
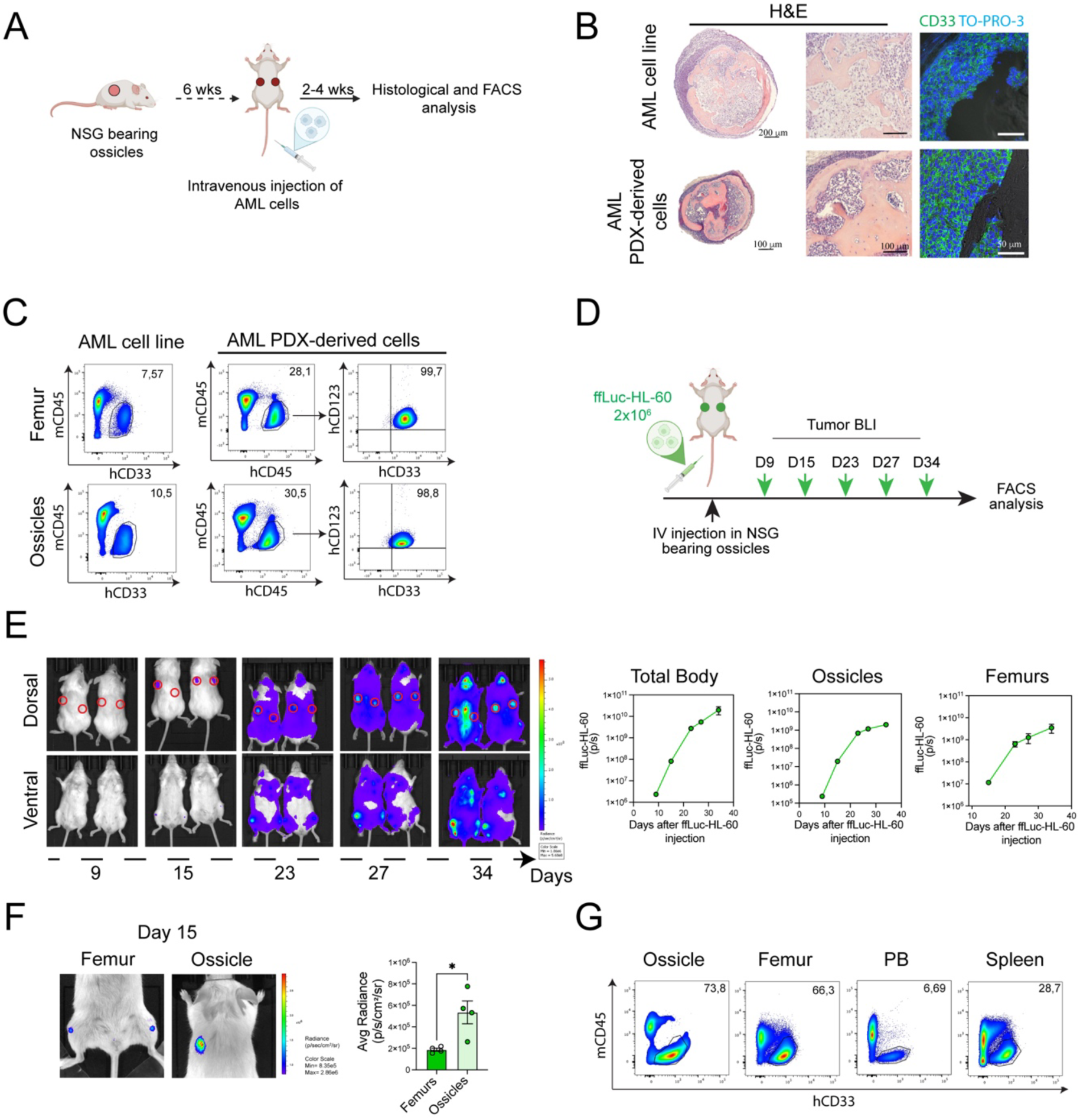
Humanized BM ossicles support AML engraftment. **A:** Experimental scheme of AML transplantation in standard ossicle-bearing NSG mice. **B:** Representative histological images of ossicles engrafted with AML cell line (HL-60, top) and pediatric AML PDX-derived cells (bottom). Sections were stained with hematoxylin and eosin (H&E) and immunostained for hCD33, demonstrating AML cell infiltration within the ossicle marrow cavity. Scale bars =200µm and 100µm (H&E) and 50µm (hCD33). Nuclei were stained with TO-PRO-3. **C:** Percentage of hCD33^+^ AML cells (KG-1) at 2-week post-transplantation and hCD45^+^/hCD33^+^/hCD123^+^ AML PDX-derived cells at 4-week post-transplantation in murine femurs and humanized ossicles. **D:** Schematic overview of the experimental setup for ffLuc-HL-60 transplantation. **E:** Left: Bioluminescence images showing the *in vivo* distribution of ffLuc-HL-60 AML cells in the total body and ossicles (red circles) from 1 to 5 weeks post-transplantation. Right: Bioluminescence quantification (total flux: photons/sec) within defined regions of interest (ROIs) to monitor leukemia burden in the total body, humanized ossicles and murine femurs (n=4 ossicles and n=4 femurs from n=2 mice). **F:** Higher magnification bioluminescence images highlighting differential engraftment rates of ffLuc-HL-60 in femurs versus ossicles at 15 days post-transplantation. Average radiance (photons/sec/cm^2^/sr) of ffLuc-HL-60 cells was compared between murine femurs and humanized ossicles. Statistical values were determined by unpaired t test. **G:** Representative percentage of hCD33^+^ cells in ffLuc-HL-60-engrafted ossicles, murine femurs, peripheral blood (PB), and spleens at the experimental endpoint.

To longitudinally assess the kinetics of AML engraftment and expansion within ossicles, we employed Firefly luciferase-labeled HL-60 cells (ffLuc-HL-60) (**Figure 1D**). Distinct bioluminescence signals were detected within the ossicles as early as two weeks post-injection (**Figure 1E**). At that time, the average radiance within the ossicles was significantly higher than that observed in the femurs, indicating that ossicles are particularly permissive to leukemic engraftment (**Figure 1F**). Over time, signal intensity increased in both humanized and murine tissues, particularly in hind limbs and spine, reflecting progressive systemic leukemia dissemination (**Figure 1E**). FACS analysis at the experimental endpoint confirmed extensive leukemic engraftment in both the ossicles and murine hematopoietic compartments, including the BM, spleen and peripheral blood (**Figure 1G**).

To evaluate the suitability of this platform for testing CAR-based immunotherapies within a species-matched microenvironment, we used cytokine-induced killer (CIK) cells engineered with a CD33.CAR via the non-viral Sleeping Beauty (SB) transposon system (*23*). Following engraftment of ffLuc-HL-60 leukemia within humanized ossicles, mice were treated with 1×10^7^ CD33.CAR-CIK cells (**Figure 2A**). CAR-CIK cells effectively trafficked to and persisted within the ossicles for up to 14 days, as demonstrated by immunohistochemistry and flow cytometry at the experimental endpoint (**Figures 2B and E**). Longitudinal BLI demonstrated that CD33.CAR-CIK treatment resulted in partial control of leukemic progression within the ossicles (**Figure 2C**). *Ex vivo* BLI of explanted ossicles further confirmed a significantly reduced leukemic signal in CAR-treated mice compared to controls (**Figure 2D**). Moreover, flow cytometric analysis of dissociated ossicles revealed a massive leukemic burden in untreated mice and a marked reduction following CD33.CAR-CIK treatment (**Figure 2E**). Collectively, these data demonstrate that humanized ossicles provide a platform in which CAR-CIK cells can engraft, persist, and exert measurable antileukemic activity within a relevant human stromal context.

**Figure 2:**
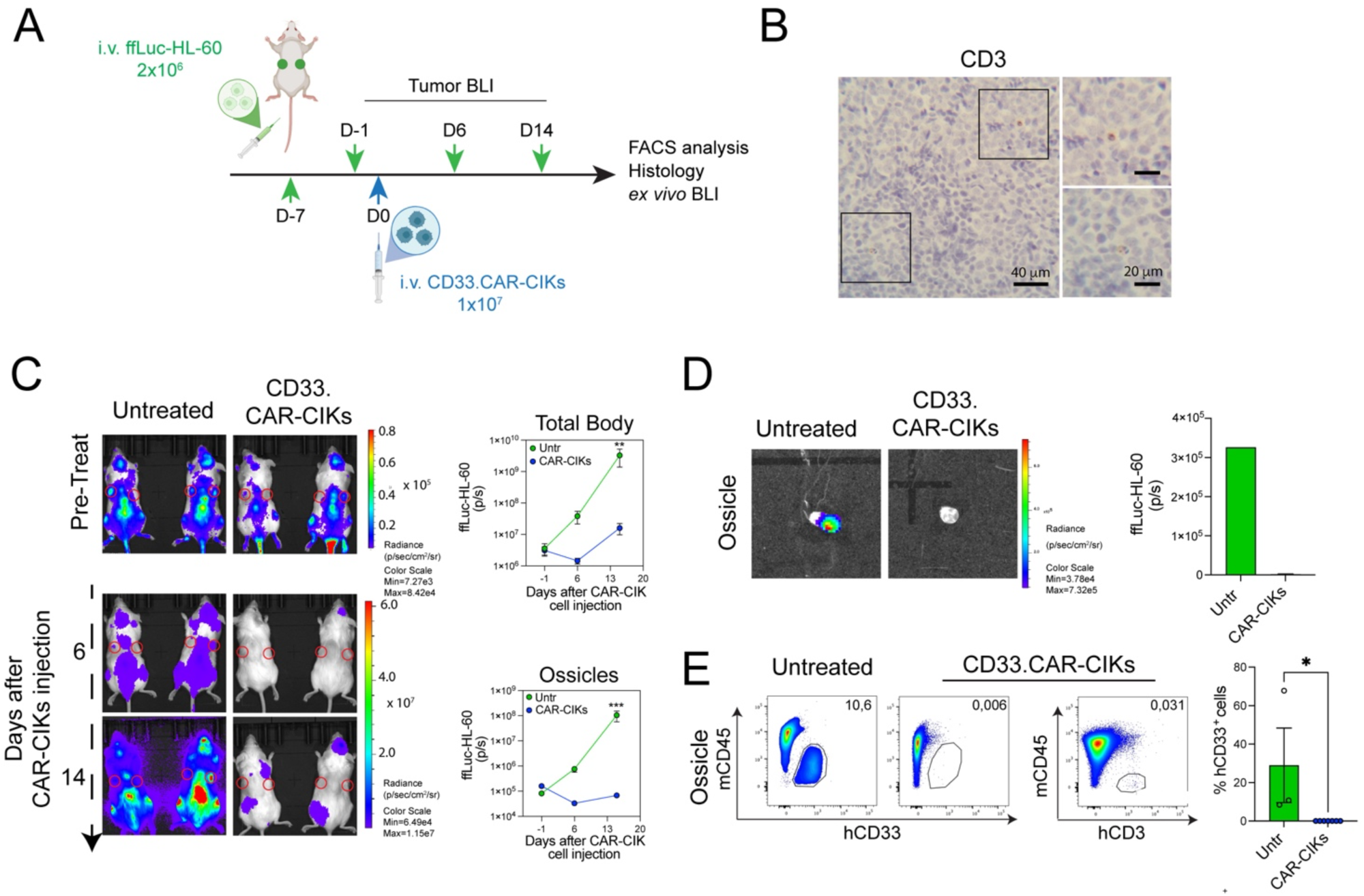
AML-engrafted humanized BM ossicles enable effective evaluation of anti-AML CAR-based immunotherapy. **A:** Experimental scheme for CD33.CAR-CIK cell treatment in AML-engrafted ossicle-bearing mice. **B:** Representative histological images of hCD3 immunostaining in explanted AML-engrafted ossicles 2 weeks after CD33.CAR-CIK cell infusion. Scale bar =40µm (left images) and 20µm (right images). **C:** Left: Bioluminescence images showing the *in vivo* distribution of ffLuc-HL-60 AML cells in the total body and ossicles (red circles) of mice, comparing untreated and CD33.CAR-CIK-treated groups. Right: Bioluminescence quantification of AML leukemia burden in the total body (top) and humanized ossicles (bottom) (n=3 untreated and n=4 treated mice). Statistical values were determined by two-way ANOVA with Sidak’s multiple comparisons test. **D:** Representative *ex vivo* bioluminescence images and corresponding quantification of ossicles derived from untreated versus CAR-CIK cell-treated mice. **E:** Left: Representative FACS dot plots showing the percentage of hCD33^+^ ffLuc-HL-60 cells and hCD3^+^ CAR-CIK cells in humanized ossicles harvested at the experimental endpoint. Right: Quantification of hCD33^+^ cells percentage in humanized ossicles from untreated versus CAR-CIK treated mice. Statistical values were determined by unpaired t test.

### Humanized BME-ossicles provide an expanded, structurally faithful BM niche that supports AML modeling

While standard cartilaginous pellet-derived ossicles support the leukemic engraftment within a well-organized human stromal niche, their reduced dimensions limit both BLI spatial resolution and cell recovery after dissociation, thereby restricting downstream analyses. To overcome these limitations, we optimized the conventional chondroid differentiation protocol by resuspending human MSCs in basement membrane extract (BME/Matrigel) (**Supplementary Figure 1A**). BME supplementation resulted in significantly larger pellets that maintained similar circular shapes and exhibited well-defined chondrogenic differentiation comparable to that of conventional cultures (**Supplementary Figures 1B-C**). More important, the presence of BME did not affect the expression of key cartilage-associated genes, including SRY-box transcription factor 9 (*SOX9)*, collagen type II *(COL2A1)*, and aggrecan *(ACAN)* (**Supplementary Figure 1D**). Consistently, both standard and BME-cartilaginous pellets showed similar immunolocalization patterns for SOX9 and COL2, whereas ACAN displayed a more focal staining pattern in BME-pellets (**Supplementary Figure 1E**).

Following *in vitro* differentiation, the capacity of BME-cartilaginous pellets to generate ossicles *in vivo* was evaluated by ectopic implantation in NSG mice. Macroscopic observation and histological analysis indicated graft vascularization and the establishment of a marrow compartment (**Figures 3A and C**). Micro-computed tomography (μCT) analysis revealed the successful formation of mature bone, characterized by well-defined cortical and trabecular structures in 3D reconstructions. Quantitative μCT analysis across multiple primary MSC populations demonstrated reproducible bone formation, with consistent bone volume to total volume (BV/TV) ratios and absolute volumes (**Figure 3B**). Notably, BME-ossicles were significantly larger than standard ossicles, providing an expanded BM cavity while maintaining a comparable marrow-to-bone ratio. Histological analyses confirmed that BME-ossicles recapitulated key anatomical features of bone, including a cortical-like bony collar and a well-organized marrow cavity containing trabecular bone and hematopoietic cells (**Figure 3C**).

**Figure 3:**
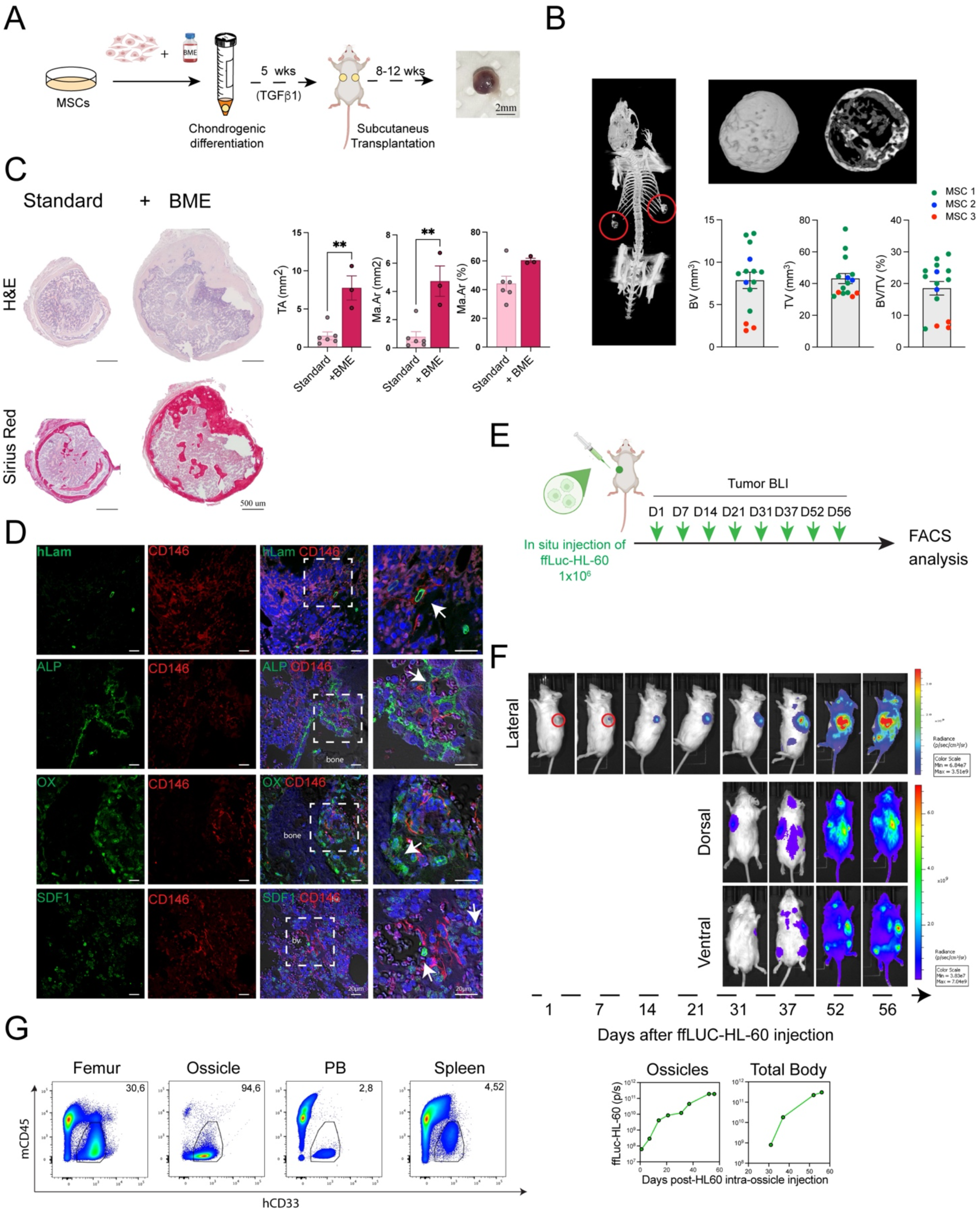
BME-ossicles recapitulate the spatial architecture and cellular composition of the human BM niche and support AML engraftment. **A:** Schematic overview of BME-ossicle generation and macroscopic appearance of explanted BME-ossicles after ectopic implantation in NSG mice. **B:** Left: 3D reconstruction of the entire skeleton of NSG mouse bearing BME-ossicles (red circles) using micro-computed tomography. Right: 3D reconstruction of a full BME-ossicle and its internal trabecular compartment (top). Morphometric analysis of BME-ossicles derived from three different MSC donors (n=15; bottom). Parameters evaluated included bone volume (BV, mm^3^), total volume (TV, mm^3^), and bone volume over total volume (BV/TV, %). Distinct colors indicate different MSC donors. **C:** Left: Representative hematoxylin and eosin and Sirius Red staining of paraffin-embedded standard and BME-ossicles. Scale bar =500µm. Right: Quantification of total area (TA, mm^2^), marrow cavity area (Ma. Ar., mm^2^) and the ratio of marrow area to total area (Ma. Ar./TA, %) in standard (n=6) versus BME-ossicles (n=3). Statistical values were determined by unpaired t test. **D:** Immunofluorescence staining of paraffin sections from BME-ossicles for human-specific LaminA/C, alkaline phosphatase (ALP), Osterix (OX, osteolineage cells), SDF1 (stromal cells) and human CD146. Nuclei were stained with Hoechst. Scale bar =20µm. bv: blood vessels. Arrows indicate double positive cells. **E:** Experimental timeline for *in vivo* tracking of ffLuc-HL-60 AML cells injected directly into the marrow cavity of BME-ossicles. **F:** Top: bioluminescence images showing the *in vivo* distribution of intra-ossicle injected ffLuc-HL-60 AML cells within BME-ossicles (red circles) and the total body. Bottom: quantification of the corresponding signal from 1 to 8 weeks post-transplantation. **G:** Representative hCD33^+^ percentages in ffLuc-HL-60-engrafted ossicles, murine femurs, peripheral blood (PB), and spleens at the experimental endpoint.

Immunofluorescence for human Lamin A/C and CD146 confirmed the presence of human-derived perivascular stromal cells, which successfully reconstituted a complex human marrow niche along with ALP- and osterix-expressing osteoblasts. Furthermore, the expression of CXCL12 (SDF1) by stromal cells underscored the functional capacity of this niche to provide essential hematopoietic support and chemotactic cues (**Figure 3D**). Consistent with ossicles established through standard protocol, the hematopoietic compartment was of recipient origin, as demonstrated by LaminA/C negativity, and included cells of erythroid (TER-119^+^), myeloid (MPO^+^), and megakaryocytic (von Willebrand factor^+^) lineages. Similarly, endothelial cells (mCD31^+^) in the vascular network were murine-derived (**Figure 3D and Supplementary Figure 2**).

Exploiting the increased volume of the BME-ossicle, we evaluated the feasibility of direct intra-ossicle delivery of ffLuc-labeled AML cells (**Figure 3E**). Serial BLI revealed rapid local engraftment and expansion, followed by leukemic egress and progressive systemic dissemination to host hematopoietic tissues (**Figure 3F**). Of note, FACS analysis at the experimental endpoint demonstrated that BME-ossicles sustain a higher leukemic burden over time compared with murine hematopoietic tissues (**Figure 3G**).

In summary, BME-ossicles provide an expanded, high-fidelity platform for AML modeling that enables precise longitudinal monitoring of leukemic progression.

### BLI-traceable CD33.CAR-CIK cells recapitulate physiological homing kinetics and maintain antileukemic activity *in vivo*

To enable the simultaneous monitoring of antileukemic activity and effector cell biodistribution, we developed traceable vectors co-expressing the CD33.CAR and a bioluminescent reporter gene. Using the non-viral Sleeping Beauty transposon system, we engineered a polycistronic construct where the reporter gene was placed downstream to the CAR transgene via a T2A self-cleaving peptide (**Figure 4A**). We evaluated two distinct reporters: Firefly Luciferase (ffLuc) and NanoLuciferase (nLuc), the latter offering a compact size for easy incorporation into complex constructs (*24*). Both constructs showed high transduction efficiency in CIK cells, as confirmed by flow cytometry showing 70-80% of CD33.CAR^+^ cells (**Figure 4B**). *In vitro* imaging demonstrated a linear correlation between CIK cell number and BLI signal intensity for both reporters, with no detectable substrate cross-reactivity (**Figure 4C**). Notably, nLuc-CD33.CAR-CIK cells produced the highest overall signal intensity. The integration of these reporters did not alter the proportions of CD4^+^ and CD8^+^ T cell subtypes and the memory profile compared to control CD33.CAR-CIK cells (**Figure 4D**). Furthermore, cytotoxicity against CD33^+^ leukemia cells and effector cytokine production showed that CIK cells engineered with either construct performed well *in vitro*, regardless of the co-expressed reporter gene (**Figure 4E**).

**Figure 4:**
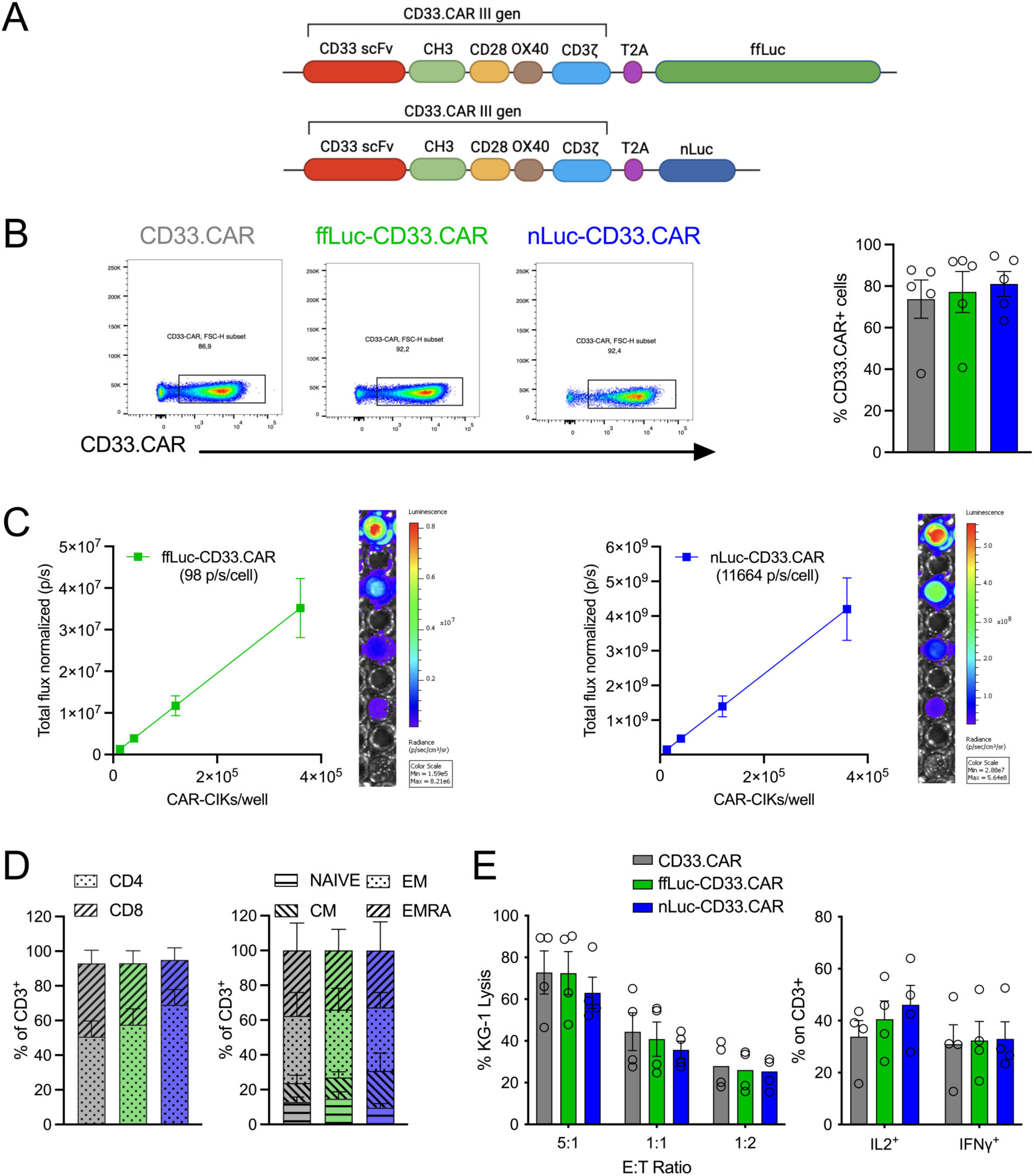
Generation and *in vitro* characterization of traceable CD33.CAR-CIK cells expressing bioluminescent reporters. **A:** Schematic of third-generation CD33.CAR constructs containing ffLuc (ffLuc-CD33.CAR) or nLuc (nLuc-CD33.CAR) as bioluminescence reporters, cloned into the pT4-transposon vector. T2A: self-cleaving peptide sequence. **B:** Representative flow cytometry plots (left panel) and corresponding quantification (right graph) showing the percentage of CIK cells transduced with standard CD33.CAR, ffLuc-CD33.CAR, or nLuc-CD33.CAR after 21 days of *in vitro* expansion (n=5 independent donors). **C:** *In vitro* BLI standard curves showing total flux (p/s) from ffLuc-CD33.CAR-CIK cells (1.3×10^4^ – 3.6×10^5^ cells) after addition of D-luciferin (left), and from nLuc-CD33.CAR-CIK cells (1.3×10^4^ – 3.6×10^5^ cells) after addition of furimazine (right) (n=5 independent donors). **D:** Quantification of CD4^+^ and CD8^+^ subsets (left) and distribution of memory subsets (right) after 21 days of *in vitro* expansion. Memory subsets include naive T cells, effector memory (EM), central memory (CM) and terminally differentiated effector memory T cells re-expressing CD45RA (TEMRA) (n=4 independent donors). **E:** Cytotoxic activity (left) and cytokine production (IL-2 and IFN-γ, right) following co-culture with CD33^+^ KG-1 AML cells across conditions after 21 days of *in vitro* expansion (n=4 independent donors).

We next characterized the *in vivo* kinetics of these traceable CARs in a conventional HL-60 AML xenograft model (**Figure 5A**). Following the intravenous infusion of 1 x 10^7^ ffLuc- or nLuc-CD33.CAR-CIK cells, we observed in both groups a rapid systemic increase in BLI signal during the initial days post-infusion, reflecting early effector cell expansion. In contrast, non-leukemic control mice exhibited a progressive reduction in signal over the same period, confirming that CAR-CIK cell proliferation is antigen-dependent. Spatiotemporal analysis of the BLI signal revealed a predominant pulmonary localization at 4 hours post-infusion (day 0.1), consistent with transient lung sequestration of infused cells, followed by progressive redistribution and homing to the femurs by 24 hours (**Figures 5B and C**). Femoral BLI signal intensity increased over time as the pulmonary signal declined, reflecting the specific accumulation and expansion of CAR-CIKs within the BM compartment, where antigen-positive leukemia cells resided. Both reporters displayed comparable trafficking and expansion kinetics (**Figure 5C**). Endpoint flow cytometry confirmed that the addition of the tracking reporters did not compromise therapeutic efficacy, as both groups achieved significant and comparable reductions in leukemic burden (**Figure 5D**).

**Figure 5:**
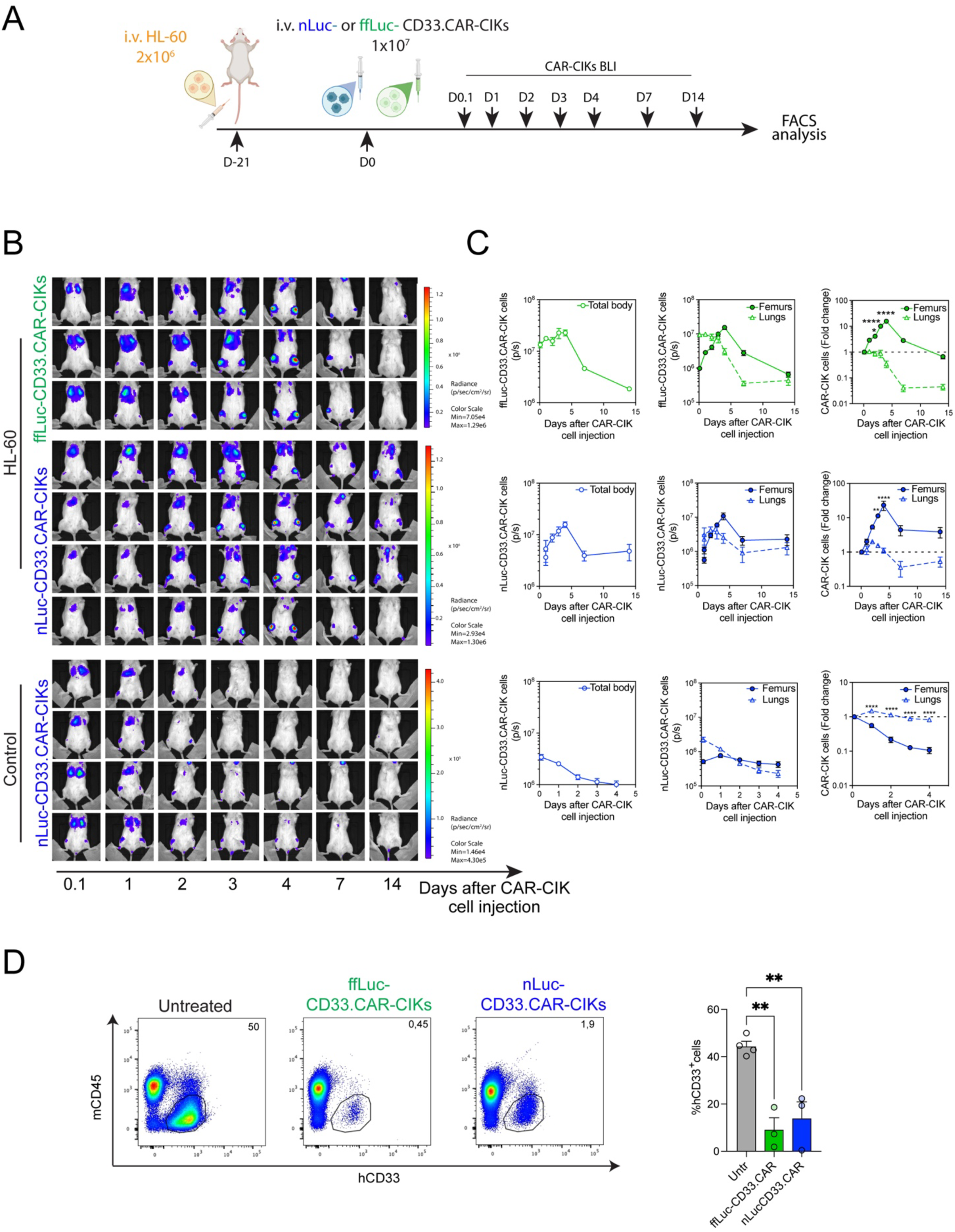
*In vivo* dynamics of traceable CD33.CAR-CIK cells in an AML xenograft model. **A:** Schematic representation of the experimental setup for *in vivo* tracking of CD33.CAR-CIK cells in AML xenografts. **B:** Bioluminescence images showing the *in vivo* distribution of intravenously administered ffLuc- and nLuc-CD33.CAR-CIK cells in HL-60-engrafted (top) or control (non-leukemic, bottom) mice at indicated time points. **C:** Bioluminescence quantification of ffLuc- and nLuc-CD33.CAR-CIK cell signals in the whole mouse body (left) and in femurs and lungs (center) across conditions at the indicated time points. The fold change of signal in femurs and lungs over time is also shown (right). Control group: n=4; nLuc-CD33.CAR-CIK: n=4; ffLuc-CD33.CAR-CIK: n=3. Statistical values were determined by two-way ANOVA with Sidak’s multiple comparisons test. **D:** Representative percentages of hCD33^+^ HL-60 cells in femurs from untreated, ffLuc- and nLuc-CD33.CAR-CIK cell-treated mice at the experimental endpoint (left). Quantification of hCD33^+^ cell percentages in murine femurs across conditions at the experimental endpoint (right). Statistical values were determined by one-way ANOVA with Sidak’s multiple comparisons test.

Collectively, these results demonstrate that the integration of BLI reporters does not perturb the biological fitness of CD33.CAR-CIKs, which exhibit the expected biodistribution kinetics and maintain antileukemic activity in AML xenograft mice.

### Dual-BLI deciphers the spatiotemporal interplay between CAR-CIKs and AML within xenograft and humanized BME-ossicle models

We next implemented a dual BLI system to longitudinally monitor the dynamic interplay between AML cells and CAR-CIK effectors *in vivo,* in both conventional xenografts and BME-ossicles. In the conventional setting, nLuc-CD33.CAR-CIK cells were infused into mice bearing established ffLuc-expressing leukemia cells (**Figure 6A**). During the initial phase of CAR-CIK expansion, leukemic progression was efficiently controlled, with precise spatial co-localisation of effector cells and leukemia within the femurs and sternum (**Figure 6C**). Specifically, leukemia control by CAR cells was associated with an early increase in the CAR-derived nLuc signal (**Figure 6B**, left panel) and a concomitant decrease in the HL60-derived ffLuc signal (**Figure 6B**, middle panel), indicating CAR cell expansion and reduction of leukemic burden, respectively. Quantitative BLI analysis confirmed the observed trend (**Figure 6B**, right upper panel). Initial leukemia control was followed by an increase in the ffLuc leukemic signal associated with a decline in the nLuc signal, indicating limited CAR-CIK persistence and highlighting the critical balance between effector persistence and leukemia relapse (**Figure 6B**, right upper panel). As expected, leukemia-associated signal remained significantly lower in CAR-treated mice compared to controls throughout the three-week study period (**Figure 6B**, right lower panel). *Ex vivo* femoral BLI and endpoint flow cytometry corroborated these findings (**Figures 6D and E**).

**Figure 6:**
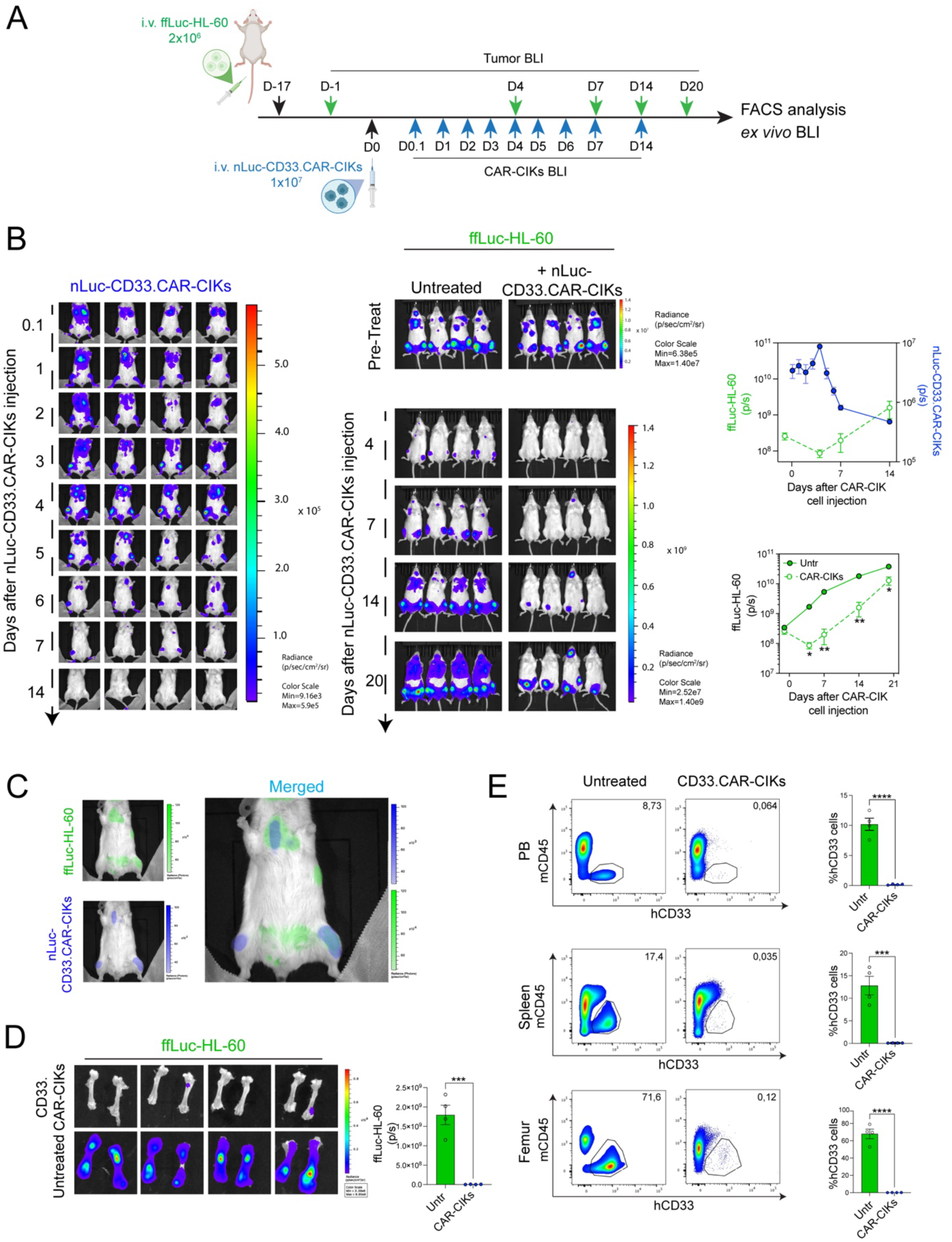
Tracking CD33.CAR-CIK cell dynamics and leukemia control by dual imaging in AML xenografts. **A:** Experimental timeline of the ffLuc-HL-60 AML xenograft mouse model treated with nLuc-CD33.CAR-CIK cells via intravenous administration. **B:** Left: BLI of nLuc-CD33.CAR-CIK cells in treated mice over time. Right: BLI of ffLuc-HL-60 leukemia in untreated and treated mice at indicated time points. Top graph: Quantification of whole-body ffLuc signal (leukemia burden) and nLuc signal (CD33.CAR-CIK cells) in treated mice, demonstrating a correlation between effector cell expansion/persistence and leukemia control. Bottom graph: Comparison of leukemia burden between untreated and treated groups (n= 4 mice per group). Statistical values were determined by two-way ANOVA with Sidak’s multiple comparisons test. **C:** Representative merged BLI images showing simultaneous detection of ffLuc-HL-60 cells (green) and nLuc-CD33.CAR-CIK cells (blue), demonstrating *in vivo* distribution and co-localization in the sternum and right femur. **D:** *Ex vivo* bioluminescence images and quantification of ffLuc-HL-60 signals in femurs from untreated versus treated mice (n= 4 mice per group; each dot represents the sum of bioluminescent signal detect in both femurs per mouse). Statistical values were determined by unpaired t test. **E:** Left: Representative percentages of hCD33^+^ HL-60 cells in PB, spleens, and femurs from untreated and nLuc-CD33.CAR-CIK cell-treated mice at the experimental endpoint. Right: Quantification of hCD33^+^ cell percentages across organs. Statistical values were determined by unpaired t test.

To evaluate these dynamics within a human microenvironment context, we combined this dual-imaging tracking strategy with AML-engrafted BME-ossicles in mice. Following direct intra-ossicle delivery of ffLuc-HL-60 cells, nLuc-CD33.CAR-CIK cells were intravenously infused once a leukemic signal had been established within the ossicle, in the absence of detectable femoral involvement. (**Figure 7A and Supplementary Figure 3**). By day 3 post-infusion, the nLuc CAR-CIK cell signal was clearly detectable within the ossicles, exhibiting a local increase of approximately 9-fold by day 7, followed by a gradual decline. In contrast, the nLuc signal in murine femurs progressively decreased over the same period, supporting antigen-dependent engagement and selective recruitment of CAR-CIK cells specifically to the engrafted ossicles (**Figures 7B**). Consistent with this preferential accumulation, CAR-treated mice exhibited significantly slower leukemic progression within the ossicles compared to untreated control animals (**Figures 7C**). Moreover, in treated mice, leukemia remained largely confined to the ossicles, whereas untreated control animals showed rapid systemic dissemination to murine femurs as early as day 7 (**Supplementary Figure 3**). Further supporting these findings, dual-BLI revealed a strict co-localisation of effector and leukemic cells within the BME ossicles (**Figure 7D**). Furthermore, total cell recovery following ossicle dissociation was significantly lower in treated mice, consistent with a reduced disease burden (**Figures 7E**), a finding supported by flow cytometric analysis demonstrating measurable antileukemic activity of CD33.CAR-CIK cells in BME ossicles harvested at sacrifice (**Figure 7F**).

**Figure 7:**
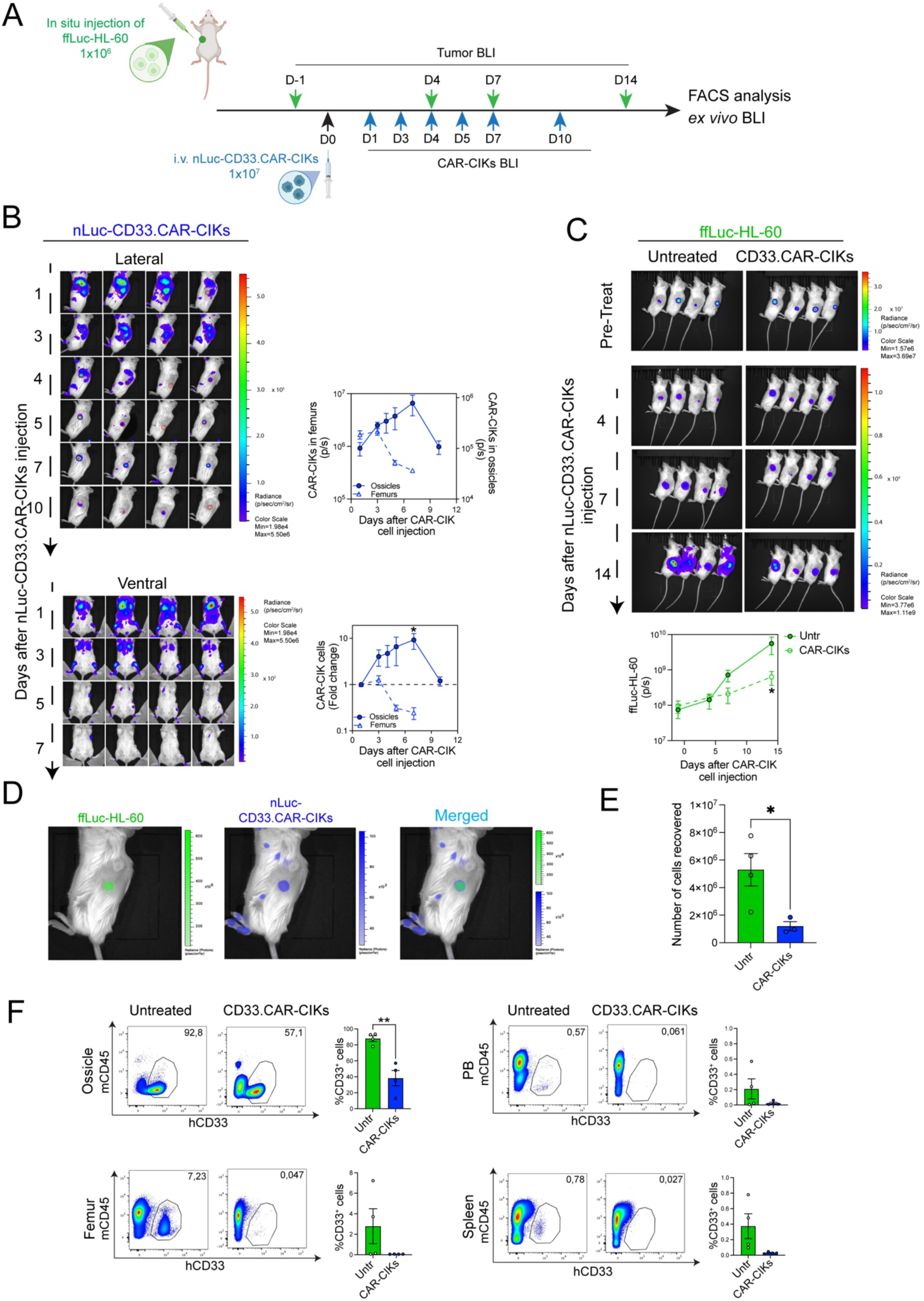
Dual BLI reveals trafficking and anti-leukemia activity of CD33.CAR-CIK cells in humanized BME-ossicles. **A:** Experimental timeline of nLuc-CD33.CAR-CIK cell treatment in mice bearing BME-ossicles. ffLuc-HL-60 AML cells were injected directly into the ossicle marrow cavity. **B:** Left: BLI in lateral view, showing the distribution of intravenously injected nLuc-CD33.CAR-CIK cells in the ossicles (top) and in ventral view, showing nLuc signals in the femurs (bottom). Right: Quantification of nLuc signal (CD33.CAR-CIK cells) in femurs and BME-ossicles of treated mice over time (top) and corresponding fold change of the nLuc signal (bottom). **C:** Top: BLI of intra-ossicle injected ffLuc-HL-60 leukemia in untreated versus treated mice at the indicated time points. Bottom: Comparison of BME-ossicles leukemia burden between untreated and treated groups (n=4 mice per group). Statistical values were determined by two-way ANOVA with Sidak’s multiple comparisons test. **D:** Representative merged BLI images showing simultaneous detection of intra-ossicle injected ffLuc-HL-60 cells (green) and intravenously injected nLuc-CD33.CAR-CIK cells (blue), demonstrating their co-localization within the BME-ossicle 7 days post-treatment. **E:** Total cell counts harvested from BME-ossicles of untreated versus treated mice at the experimental endpoint. Statistical values were determined by unpaired t test. **F:** Representative FACS dot plots and quantification of the percentage of hCD33^+^ HL-60 cells in BME-ossicles, murine femurs, PB, and spleens from untreated and nLuc-CD33.CAR-CIK cell-treated mice at the experimental endpoint. Statistical values were determined by unpaired t test.

Together, these data demonstrate that the humanized BME-ossicle platform, integrated with dual-BLI, enables high-resolution, spatially resolved monitoring of the different kinetics between AML and CAR-CIK cells.

## Discussion

Advances in immunotherapeutic approaches for AML have highlighted the critical role of the BM microenvironment as a key determinant of therapeutic efficacy, underscoring how leukemia–niche interactions can profoundly influence treatment response and resistance (*3*, *25–27*). This has emphasized the urgent need to further refine experimental platforms capable of faithfully modeling these complex interactions and enabling the evaluation of next-generation CAR-based strategies specifically designed to overcome microenvironment-mediated barriers.

Although conventional xenograft models have substantially contributed to our understanding of leukemia progression and treatment response, their reliance on a murine marrow microenvironment limits their ability to accurately recapitulate the complex interactions among leukemic cells, therapeutic effectors, and the human BM niche (*4*, *28*).

Our study addresses this unmet need by establishing a humanized ossicle-based AML platform that supports robust engraftment of leukemic blasts within a human BM microenvironment, while simultaneously allowing longitudinal assessment of CAR therapy. To our knowledge, this study provides the first direct evidence that humanized ossicles constitute a suitable experimental model for the rigorous evaluation of AML engraftment dynamics in parallel with longitudinal CAR-based immunotherapy responses within a human stromal context.

Our findings confirm that ossicles generated from primary human MSCs support both AML cell line and patient-derived xenograft cell engraftment with efficiency comparable to or exceeding that of native murine BM (*16*). Both intravenous and intra-ossicle injection strategies produced reproducible disease that mimics key histopathological features of human AML, including extensive infiltration of myeloid blasts within a well-organized marrow cavity containing human-derived perivascular stromal cells expressing CXCL12. The relevance of this model is further supported by the capacity of CD33.CAR-CIK cells to traffic to, persist within, and exert measurable antileukemic activity in the ossicles. However, the relatively small size of standard ossicles limits both the spatial resolution of bioluminescence signal and the cell recovery after ossicle dissociation.

Considering this key limitation, an important advance in our work is the technical refinement of the ossicle generation protocol through BME supplementation, which significantly increases ossicle size and marrow cavity volume while preserving overall structural characteristics. BME-supplementation in the chondroid differentiation protocol resulted in significantly larger pellets that maintain well-defined chondrogenic differentiation required for ossicle formation. In fact, the derived BME-ossicles preserve the architectural complexity of bone and BM essential for translational relevance, including a cortical-like bony collar, trabecular bone structures, and a well-organised hematopoietic compartment, containing erythroid, myeloid, and megakaryocytic lineages. Moreover, the preservation of key niche-associated elements, including CD146 positive stromal cells, osteoblastic populations and CXCL12-expressing supportive cells, indicate that ossicle enlargement does not compromise biological fidelity (*29*). Importantly, the enlarged volume of BME-ossicles enables direct intra-ossicle injection of AML cells, allowing precise local engraftment with defined cell numbers. This approach overcomes the variability associated with systemic administration, where engraftment sites and timing are less controllable, thereby improving the standardization of preclinical experiments and facilitating dose-response studies. In addition, the increased marrow cavity volume facilitates cell recovery for downstream analyses and improves the spatial resolution of *in vivo* imaging, thereby increasing the overall efficiency and experimental versatility of the platform.

An additional strength of this platform is its use of primary human MSCs to generate the ossicles. Unlike models based on engineered or immortalized stromal cell lines, this approach creates an important future application to establish fully autologous leukemic niches by combining patient-derived MSCs and leukemic cells, thereby enabling the investigation of leukemia–microenvironment interactions and immunotherapeutic responses in a patient-specific setting (*30*, *31*).

A further key aspect of this work is the integration of CAR cell therapy with the humanized ossicle-based model, which enables the functional evaluation of CAR activity within a physiologically faithful human BM microenvironment. Building on this framework, we developed and validated a dual BLI system using distinct luciferases (ffLuc for AML, and nLuc for CAR-CIK) which enables simultaneous, non-invasive longitudinal visualization of both leukemic burden and CAR-CIK cell dynamics within the same animal. Although BLI has become a standard tool for tracking tumor burden or adoptively transferred immune cells individually, the simultaneous monitoring of both populations within a humanized BM niche provides a substantially more informative framework for studying therapeutic responses. Reporter integration in our CAR transposon platform demonstrated high transduction efficiency without compromising CAR-CIK biological fitness, preserving memory phenotype and effector function. *In vivo*, traceable CAR-CIK cells exhibited antigen-dependent expansion kinetics and a physiologically relevant biodistribution pattern, characterised by rapid pulmonary sequestration followed by homing to the leukemic BM (*18*). Importantly, application of the dual BLI system revealed details about the spatiotemporal interplay between CAR-CIK cells and AML blasts, confirming the critical balance between effector persistence and leukemia control (*32*). CAR-CIK expansion closely correlated with leukemia control, whereas contraction of the effector population preceded disease progression.

Application of dual-BLI to humanized BME-ossicles revealed antigen-dependent selective recruitment of CAR-CIK cells specifically within the leukemic ossicles. The strict co-localization of effectors and leukemic cells and significantly slower leukemic progression in CAR-treated mice demonstrate that this platform enables high resolution monitoring of therapeutic dynamics within the physiologically relevant human microenvironment.

While our humanized BME-ossicle platform represents a significant advance, several limitations should be acknowledged. Although the stromal compartment is of human origin, the endothelial and hematopoietic components remain predominantly murine, and the model lacks a fully functional human immune system (*33*). Consequently, some aspects of immune regulation and therapy response cannot yet be fully recapitulated. Future efforts integrating human immune compartments into the ossicle platform may further increase its translational relevance (*15*, *16*, *34*). The dual-BLI approach requires genetic modifications of both AML and CAR-engineered cells, which may limit applicability to some primary patient samples that cannot be efficiently transduced.

Beyond the evaluation of CD33.CAR-CIK cells, this platform may be widely applicable to the preclinical assessment of next-generation cellular therapies, including armored CARs and combinatorial immunotherapeutic strategies designed to overcome microenvironment-mediated resistance mechanisms (*35*). In particular, the dual-tracking approach could facilitate the evaluation of engineering strategies aimed at enhancing CAR-cell trafficking, persistence, and functionality within the BM niche (*36*). For example, our previous work demonstrated that enforced CXCR4 expression significantly enhanced CAR-CIK cell homing to the CXCL12-rich BM microenvironment and ameliorated AML control, highlighting the importance of niche-directed trafficking for therapeutic efficacy (*37*). Leveraging the present model, simultaneous monitoring of leukemic burden and effector-cell dynamics could provide direct insight into whether such benefits derive from increased recruitment, prolonged persistence, enhanced local expansion, or a combination of these mechanisms. Likewise, this system could be exploited to evaluate armored CARs engineered to express different immune-stimulatory cytokines (e.g., IL-15, IL-18, or IL-7/CCL19), as well as strategies aimed at overcoming stromal-mediated immunosuppression through the modulation of MSCs activity, disruption of leukemia–stroma interactions, or targeting of suppressive pathways (*1*, *9*). By coupling longitudinal imaging with multiparametric analyses, including flow cytometry, single-cell RNA sequencing, and spatial transcriptomics, this platform may provide a comprehensive framework for dissecting CAR-cell behavior, therapeutic efficacy, and mechanisms of resistance within a humanized BM niche.

In conclusion, our findings establish humanized ossicles as a robust and versatile platform for modeling AML and evaluating CAR-based immunotherapies within a bona fide human BM microenvironment. The generation of enlarged ossicles suitable for controlled local leukemic inoculation, combined with the implementation of dual BLI, substantially expands the experimental capabilities of this system by enabling the simultaneous longitudinal monitoring of leukemic burden and effector-cell dynamics *in vivo*. By bridging the gap between conventional xenograft models and the complexity of the human marrow niche, this platform offers the unique opportunity to investigate niche-dependent mechanisms governing leukemia progression, immune-cell trafficking, therapeutic response, and resistance. We anticipate that these advances will facilitate the development and optimization of next-generation CAR-based strategies and accelerate their translation toward more effective therapies for AML.

## Material and Methods

### AML cell lines and PDX cells

KG-1, HL-60, and ffLuc-HL-60 AML cell lines were obtained from the American Type Culture Collection (ATCC). KG-1 cells were maintained at a density of 0.3×10^6^ cells/ml in complete medium consisting of RPMI 1640 (Sigma-Aldrich) supplemented with 10% heat-inactivated fetal bovine serum (FBS, Gibco), 2 mM L-glutamine, 25 IU/ml penicillin, and 25 mg/ml streptomycin (Lonza). HL-60 and ffLuc-HL-60 cell lines were maintained at 0.6×10^6^ cells/ml in complete medium supplemented with 20% FBS.

Pediatric AML patient-derived xenograft (PDX) cells were harvested from the bone marrow (BM) or spleen of engrafted NSG mice and cryopreserved until required for *in vivo* injection.

### MSC isolation and culture

Mononuclear cells (MNCs) were isolated from healthy donor BM samples via Ficoll-Paque (Cytiva) density gradient centrifugation. MNCs were seeded at a density of 2×10^5^ cells/cm^2^ in complete Minimum Essential Medium alpha (MEM α, Sigma-Aldrich) supplemented with 20% heat-inactivated FBS, 2 mM L-glutamine, 25 IU/ml penicillin, and 25 mg/ml streptomycin. Cells were maintained at 37°C in a humidified atmosphere containing 5% CO_2_. Non-adherent cells were removed 24 hours after initial plating to enrich MSCs via plastic adherence. The culture medium was changed twice a week. Upon reaching 70-80% confluence, MSCs were detached by trypsin, replated, and expanded for a maximum of 4 passages before cryopreservation for subsequent experiments. The study was approved by the internal Ethics Committee and all procedures were performed following relevant guidelines and regulations.

### Generation of standard and BME-cartilaginous pellets

#### Standard cartilaginous pellets

To generate standard cartilaginous pellets from MSCs, a micro-mass culture system was used as previously described (*13*), with minor modifications. Briefly, after expansion in growth medium, MSCs were harvested using trypsin (Euroclone) and transferred into 15 ml polypropylene conical tubes at a density of 3×10^5^cells/tube. Cells were centrifuged at 1500 rpm for 5 minutes and cultured for 3-5 weeks in chondrogenic differentiation medium (CDM) consisting of Dulbecco’s Modified Eagle Medium high glucose (DMEM, Gibco) supplemented with 0.2% bovine serum albumin (BSA, Sigma-Aldrich), ITS^TM^ Premix (BD Biosciences), 1 mM sodium pyruvate (Gibco), 50 μg/ml 2-phosphate-ascorbic acid (Sigma-Aldrich), 100 nM dexamethasone (Sigma-Aldrich), 0.1 mM non-essential amino acid solution (Gibco), and 10 ng/ml transforming growth factor (TGF)-β1 (Sigma-Aldrich).

#### BME-cartilaginous pellets

To generate BME-cartilaginous pellets, 3×10^5^ MSCs were resuspended in 50 μl of basement membrane matrix (BME/Matrigel high concentration, Corning). The cell/BME mixture was allowed to gel at 37°C for 10 minutes before CDM addition and cultured for 4-5 weeks in CDM as reported above.

### RNA extraction and real-time PCR

Gene expression analysis was performed on standard and BME-cartilaginous pellets. Total RNA was extracted using TRIzol reagent (Invitrogen), following the manufacturer’s protocol. For each sample, 1μg of total RNA was reversed transcribed using SuperScript^TM^ II Reverse Transcriptase (Invitrogen) in the presence of random hexamers (Invitrogen). Real-time PCR was performed on a 7500 Fast Real-Time PCR System (Applied Biosystem) using SYBR Green Master Mix (Thermo Fisher Scientific) and the specific primers listed in **Supplementary Table 1**. Glyceraldehyde 3-phosphate dehydrogenase (GAPDH) was used as the housekeeping gene for reference. Relative gene expression was quantified using the comparative threshold cycle (Ct) method (2^-DDCt^).

## Histology, histochemistry, and histomorphometry

Cartilage pellets and ossicles were fixed in 4% buffered formaldehyde solution for 12 hours at 4°C. Following fixation, ossicles were decalcified in 10% EDTA prior to paraffin embedding according to standard procedures. Sections (4-μm-thick) were stained with hematoxylin and eosin (H/E) for general morphological analysis or with Toluidine Blue to evaluate the cartilaginous matrix.

For histomorphometric analysis, the area and circularity of the cartilage pellets and the total area and the fraction occupied by hematopoietic tissue in the ossicles were quantified using ImageJ software.

### Immunohistochemistry and immunofluorescence staining

For immunohistochemistry, 4-μm-thick paraffin-embedded sections of cartilage pellets or ossicles were immunostained with the following primary antibodies: SOX9 (1:1000, ab185966, Abcam), Aggrecan (1:100, ab3778, Abcam), Collagen type II (1:200, ab53047, Abcam), von Willebrand factor (A0082, Dako), MPO (myeloperoxidase, 760-2659, Roche), TER-119 (550565, BD Biosciences), and CD3 (1:200, A0452, Dako). When necessary, antigen retrieval was performed using heat-mediated or enzymatic methods prior to primary antibody incubation. Endogenous peroxidases were quenched by incubating slides in 3% hydrogen peroxide (Carlo Erba) for 40 minutes. Non-specific binding was blocked with 10% BSA in PBS. Following primary antibody incubation, slides were repeatedly washed in PBS and then were incubated with horseradish peroxidase (HRP)-conjugated goat anti-rabbit IgG (31460, Thermo Fisher Scientific) or HRP-conjugated goat anti-mouse IgG (31430, Thermo Fisher Scientific) for 30 minutes at room temperature in a humidified chamber. The peroxidase reaction was developed using 3,3’-diaminobenzidine (DAB) substrate kit (SK-4105, Vector Laboratories).

For immunofluorescence, 4-μm-thick paraffin sections were immunostained with following primary antibodies: human Lamin A/C (1:500, EPR4100, Abcam), CD146 (1:40, AF932, R&D Systems), CD33 (1:100, ab269456, Abcam), Osterix (1:200, ab22552, Abcam), SDF-1 (1:100, ab25117, Abcam), CD31 (1:100, ab182981, Abcam), and ALP (1:100, 11187-1-AP, Proteintech). Heat-induced antigen retrieval and blocking with 5% BSA in PBS were performed before antibody incubation. After washing in PBS, sections were then incubated with Alexa Fluor 488-conjugated donkey anti-rabbit IgG (A-21206, Invitrogen) or Alexa Fluor 594-conjugated donkey anti-goat IgG (A-32758, Invitrogen) for 30 minutes at room temperature. Nuclei were counterstained with Hoechst 33342 (62249, Thermo Fisher Scientific). Sections were mounted using Vectashield anti-fade mounting medium (H-1000, Vector Laboratories) and images were acquired using a Zeiss LSM 710 confocal laser scanning microscope (Zeiss).

### Traceable CAR transposon design

Two bicistronic Sleeping Beauty (SB) transposon vectors, CD33.CAR-T2A-ffLuc and CD33.CAR-T2A-nLuc, were generated by modifying a parental SB pT4-transposon vector containing a CD33-specific third-generation CAR construct (*37*). Each cassette comprised a cDNA sequence encoding the CAR followed by a self-cleaving T2A peptide in frame with an imaging reporter gene. The bioluminescence reporters used were firefly luciferase (ffLuc) or nano-luciferase (nLuc) (*24*). The SB-pT4 vector and SB100X transposase were kindly provided by Dr. Izsvak (Max-Delbruck-Center for Molecular Medicine, Berlin, Germany).

### Preparation of CAR-CIK cells

CIK cells were generated, as previously described (*23*), starting from human peripheral blood mononuclear cells (PBMCs) isolated from healthy donors using Ficoll-Paque density gradient centrifugation. On day 0, PBMCs were seeded at 3×10^6^ cells/ml in Advanced RPMI medium supplemented with 10% FBS, 2 mM L-glutamine, 25 IU/ml of penicillin, 25mg/ml of streptomycin, and interferon-gamma (IFN-γ; 1000 U/ml; Dompe Biotec). On day 1, interleukin-2 (IL-2; 300 U/ml; ‘Chiron BV) and anti-CD3 monoclonal antibody OKT3 (50 ng/ ml; Janssen-Cilag) were added. For non-viral CAR engineering, activated PBMCs were nucleofected with 15 μg of supercoiled DNA transposon plasmid (encoding either monocistronic CD33.CAR or bicistronic constructs) and 0.5 μg of supercoiled DNA encoding the SB100x transposase. Nucleofection was performed using the Amaxa 4D-NucleofectorTM (Lonza) with the P3 Primary Cell 4D-Nucleofector kit (Lonza), applying the pre-set EO-115 program. Subsequently, fresh complete medium and IL-2 were added twice a week during 21 days of culture. Cell concentration was maintained at 1×10^6^ cells/ml throughout expansion. At the end of the differentiation protocol, CIK cells consisted predominantly of CD3^+^ T lymphocytes expressing CD8 and CD56 and exhibited an effector memory phenotype.

### Immunophenotypic characterization of CAR-CIK cells

CD33.CAR expression on CIK cells was detected via a sequential staining protocol. Briefly, cells were incubated with a recombinant human sialic acid binding Ig-Like Lectin 3/Siglec-3/CD33 protein with a Fc and a 6xHis tag at the C-terminus (C-Fc-6His) (Gentaur), followed by an Alexa Fluor 647-conjugated anti-human IgG Fc secondary antibody (LiStarFish).

Immunophenotyping for CD4 and CD8 expression and T cell memory subsets was performed using fluorochrome-conjugated antibodies against CD3 (clone SK7), CD8 (clone RPA-T8), CD4 (clone SK3) (BioLegend), CD56 (clone B159), CD62L (clone DREG-56), CD45RO (clone UCHL1) (BD Biosciences).

Samples were acquired using a FACSCanto II flow cytometer (BD Biosciences) at the end of 21 days of culture and analyzed using FlowJo v10.10.0 software (Tree Star Inc.).

### CAR-CIK functional assays

#### Cytotoxicity assay

Target CD33^+^ KG-1 AML cells were labeled with CellTracker Violet BMQC (Invitrogen) and co-cultured with either control or luciferase-labeled CD33.CAR-CIK cells for 4 hours. Assays were performed at effector-to-target (E:T) ratios of 5:1, 1:1, and 1:2 in 96-well round-bottom plates. Target cell mortality was quantified by flow cytometry using the Annexin V Apoptosis Detection Kit with 7-AAD (Biolegend) following the manufacturer’s instructions. Analysis was performed by gating on the Violet^+^ target cell population.

#### Intracellular Cytokine Staining (ICS) assay

Control or luciferase-labeled CD33.CAR-CIK cells were co-cultured with KG-1 cells for 5 hours at an E:T ratio of 1:3 in 96-well round-bottom plates. After 2.5 hours of co-culture, BD GolgiStopTM was added to inhibit cytokine secretion, and incubation was continued for an additional 2.5 hours. Cells were first stained for CD3 surface expression, followed by fixation and permeabilization using the BD Cytofix/Cytoperm kit for intracellular staining with anti-human IFN-γ (clone B27) and IL-2 (clone MQ1-17H129) mAbs (BD Biosciences). Samples were acquired by flow cytometry and analyzed by to inhibit cytokine secretion, gating on the CD3^+^ population.

### *In vitro* quantification of bioluminescent signal

To evaluate the correlation between bioluminescent intensity and CAR-CIK cell number, serial dilutions of nLuc- or ffLuc-CD33.CAR-CIK cells were seeded in duplicate in 96-well black flat-bottom plates in 200 μl of culture medium. Cells were allowed to settle for at least 30 minutes at 37°C. Subsequently, D-Luciferin potassium salt (ffLuc substrate; 150 µg/ml final concentration; Revvity) in PBS or furimazine (nLuc substrate; 2 µg/ml final concentration; Tebubio) in DMSO was added to the corresponding wells. Bioluminescent signals were captured using the IVIS Lumina III imaging system (Perkin Elmer) with an open filter. Light output was quantified by drawing regions of interest (ROIs) around each well, and the total flux in photons per second (p/s) was calculated using Living Image 4.8.2 Software (Revvity). A background control (medium only) was included in all *in vitro* assays.

### In vivo studies

All animal experiments were conducted in accordance with national and international guidelines and approved by the Italian Ministry of Health (Authorization No. 178/2023-PR). All procedures were performed following protocols approved by the Institutional Animal Care and Use Committee (Organismo Preposto al Benessere Animale, OPBA) of the University of Milano-Bicocca. Six- to eight-week-old female NSG (NOD.Cg-*Prkdc^scid^Il2rg^tm1Wjl^*/SzJ) mice were purchased from Charles River Laboratories.

### Subcutaneous implantation of cartilaginous pellets

Standard or BME-cartilaginous pellets were implanted on the back of NSG mice (1-2 pellets per animal) under aseptic surgery conditions, as previously described (*13*). Briefly, the dorsal area was shaved and subcutaneous pouches were created and filled with pellets. Surgery was performed under anesthesia (isoflurane 4% induction, 2% maintenance). Non-absorbable sutures were used to close the wounds.

### *In vivo* μCT scanning

Upon successful formation of the BME ossicles, mice underwent *in vivo* micro-computed tomography (μCT) imaging. Animals were anesthetized with isoflurane and positioned in the prone position. Scans of the lower body were acquired using a SkyScan 1176 micro-CT system (Bruker) equipped with a tungsten X-ray source. The scanning parameters were set at 50 kV, 500 μA, 18 μm voxel size, and 210 ms exposure time. A 180° rotation scan was performed with a rotation step of 0.5°. Image reconstruction was carried out using NRecon software (Bruker). Quantitative analysis of total volume (TV) and bone volume (BV) was performed using ImageJ software with the BoneJ plugin by measuring the total tissue volume and the volume of highly mineralized tissue.

### *In vivo* AML transplantation and treatment models

Six weeks post-implantation of standard pellets or upon successful BME-ossicle maturation, mice were transplanted with AML cells via either intra-ossicle or intravenous injections.

#### Intra-ossicle injections

Mice were anesthetized (isoflurane 4% for induction, 2% for maintenance) and the skin was shaved. Each BME-ossicle was held using forceps and 1×10^6^ AML cells in 15 μl of PBS were injected directly into the cavity using a 30G needle.

#### Intravenous injections

Mice were immobilized in a plastic restrainer and 2×10^6^ AML cells in 200 μl PBS were injected into the tail vein.

For treatment with CAR-CIK cells, mice received 1×10^7^ CD33.CAR-CIK, ffLuc-CD33.CAR-CIK, or nLuc-CD33.CAR-CIK cells via tail vein injection.

Leukemia burden and CAR-CIK cell distribution were monitored using BLI at multiple time points, as described below.

### In vivo BLI

The distribution of ffLuc-HL-60 cells or ffLuc-CD33.CAR-CIK cells was analyzed following intraperitoneal injection of D-Luciferin (150 mg/kg body weight, approximately 200 μL/mouse). Isoflurane-anesthetized mice were imaged using the IVIS Lumina III system with an open filter, 5 minutes post-substrate injection.

The trafficking of nLuc-CD33.CAR-CIKs was monitored at the indicated timepoints, from 4 hours to 14 days post-administration. Furimazine was administered to anesthetized mice via intravenous injection (100 μg/ml, 100 μl/mouse), and animals were imaged immediately. Images of mice were acquired in dorsal, ventral, or lateral positions. BLI data were analyzed using Living Image software. Unless otherwise stated, the BLI signal is reported as total flux (photons/second).

### Simultaneous *in vivo* BLI

To achieve simultaneous capture of both luciferase signals, mice underwent tail vein cannulation. Following isoflurane anesthesia, furimazine was administered intravenously via the cannula and mice were immediately imaged using the IVIS Lumina III to capture the nLuc-CD33.CAR-CIK signal. Following a 30-minutes washout period to allow furimazine clearance, the cannula was flushed to remove any residual substrate. D-luciferin was then administered intravenously and the bioluminescent signal was acquired to detect ffLuc-HL-60 cells. The signal from each cell type was displayed using pseudocolors and overlaid onto a photographic image of the mouse to identify anatomical distribution and the co-localization of the two distinct cell populations.

### Ex vivo BLI

At the experimental endpoint, mice were administered with D-luciferin intraperitoneally and sacrificed after 5 minutes. Leukemia-engrafted femurs and BME-ossicles were harvested, placed in Petri dishes, and imaged using the IVIS Lumina III with open filters to quantify the ffLuc signal.

### Flow cytometry

At the experimental timepoint, mice were euthanized and the femurs, spleens, peripheral blood, and ossicles were harvested and processed into single-cell suspensions. Femurs and ossicles were flushed with 1x PBS with 1% FBS. Spleens were gently dissociated using the flat end of a syringe plunger. All organs underwent red blood cell lysis using Ammonium-Chloride-Potassium (ACK) lysing buffer (Voden Medical), followed by two washes with 1x PBS with 1% FBS. Cells were counted using Trypan blue (Sigma-Aldrich) exclusion or an automated cell counter. Subsequently, cells were stained in 100 μl of 1x PBS + 1% FBS using the following antibodies: anti-mCD45-PE (1:3500, clone 30-F11, Thermo Fisher Scientific), anti-hCD33-PE-Cy7 (1:30, clone P67.6, BD Biosciences), anti-hCD3-PerCP (1:100, clone SK7, BioLegend), anti-hCD45-Pacific Orange (1:100, clone HI30, Thermo Fisher Scientific). Cells were incubated for 15 min at room temperature before washing. Samples were acquired on a FACS Canto II instrument (BD Biosciences) and data was analyzed using FlowJo software (version 10.10.0).

## Statistical analyses

Unless otherwise stated, data are presented as mean ± standard error of the mean (SEM). Statistical differences between two groups were analyzed using Student’s t test. For longitudinal and repeated-measures analyses, either a mixed-effects model or repeated-measures ANOVA was used, depending on variance and distribution characteristics. The Greenhouse-Geisser correction was applied to account for violations of sphericity, and multiple pairwise comparisons were adjusted using the Holm-Sidak method. Statistical tests and p values are indicated in the respective figure legend, with * p<0.05, ** p<0.01, *** p<0.005. All graphs and statistical analyses were performed using GraphPad Prism (v10.2.3). Illustrations were created by BioRender.com.

## Declaration of generative AI in the manuscript preparation process

During the preparation of this work the authors used ChatGPT in order to polish the language. After using this tool/service, the authors reviewed and edited the content as needed and take full responsibility for the content of the published article.

## Acknowledgments

The authors acknowledge Euro-BioImaging (www.eurobioimaging.eu) for providing access to imaging technologies and services via the Multi-Modal Molecular Imaging Italian Node (IBSBC-CNR, Segrate, Italy).

## Author contributions

Conceptualization: M.S., and A.P.; methodology: S.D., M.A., A.P., M.B., E.R. and S.T.; investigation: S.D., M.A., V.Z., C.G., E.G., G.A. and A.P.; visualization: S.D., M.A. and V.Z.; writing – original draft: A.P. and S.D.; writing – review and editing: M.S., M.R. and A.B.; funding acquisition: M.S.; supervision: M.S. and A.P.

## Data availability

All data needed to evaluate the results in the paper are present in the paper and/or the Supplementary Materials. Requests for further information, resources, and reagents should be directed to and will be fulfilled by the corresponding author, Prof. Marta Serafini.

## References

1. M. Khalifeh, E. Hopewell, H. Salman, CAR-T cell therapy for treatment of acute myeloid leukemia, advances and outcomes. Molecular Therapy 33, 2441–2453 (2025).

2. Honing CAR T cells to tackle acute myeloid leukemia | Blood | American Society of Hematology. https://ashpublications.org/blood/article/145/11/1113/534311/Honing-CAR-T-cells-to-tackle-acute-myeloid.

3. S. Tettamanti, A. Pievani, A. Biondi, G. Dotti, M. Serafini, Catch me if you can: how AML and its niche escape immunotherapy. Leukemia 36, 13–22 (2022).

4. A. Dozzo, A. Galvin, J.-W. Shin, S. Scalia, C. M. O’Driscoll, K. B. Ryan, Modelling acute myeloid leukemia (AML): What’s new? A transition from the classical to the modern. Drug Deliv Transl Res 13, 2110–2141 (2023).

5. B. B. Duncan, C. E. Dunbar, K. Ishii, Applying a clinical lens to animal models of CAR-T cell therapies. Molecular Therapy - Methods & Clinical Development 27, 17–31 (2022).

6. N. Baryawno, D. Przybylski, M. S. Kowalczyk, Y. Kfoury, N. Severe, K. Gustafsson, K. D. Kokkaliaris, F. Mercier, M. Tabaka, M. Hofree, D. Dionne, A. Papazian, D. Lee, O. Ashenberg, A. Subramanian, E. D. Vaishnav, O. Rozenblatt-Rosen, A. Regev, D. T. Scadden, A Cellular Taxonomy of the Bone Marrow Stroma in Homeostasis and Leukemia. Cell 177, 1915–1932.e16 (2019).

7. K. Aoki, M. Kurashige, M. Ichii, K. Higaki, T. Sugiyama, T. Kaito, W. Ando, N. Sugano, T. Sakai, H. Shibayama, H. C. B. Club, A. Takaori-Kondo, E. Morii, Y. Kanakura, T. Nagasawa, Identification of CXCL12-abundant reticular cells in human adult bone marrow. British Journal of Haematology 193, 659–668 (2021).

8. S. Méndez-Ferrer, D. Bonnet, D. P. Steensma, R. P. Hasserjian, I. M. Ghobrial, J. G. Gribben, M. Andreeff, D. S. Krause, Bone marrow niches in haematological malignancies. Nat Rev Cancer 20, 285–298 (2020).

9. A. Pievani, M. Biondi, S. Tettamanti, A. Biondi, G. Dotti, M. Serafini, CARs are sharpening their weapons. Journal for ImmunoTherapy of Cancer 12 (2024).

10. I. Zugasti, Lady Espinosa-Aroca, K. Fidyt, V. Mulens-Arias, M. Diaz-Beya, M. Juan, Á. Urbano-Ispizua, J. Esteve, T. Velasco-Hernandez, P. Menéndez, CAR-T cell therapy for cancer: current challenges and future directions. Sig Transduct Target Ther 10, 210 (2025).

11. K. Prasad, R. S. Cross, M. R. Jenkins, Progress in the development of cytokine armoured CAR T cells. Nat Rev Immunol, 1–13 (2026).

12. W. den Hartog, J. Harwood, S. Kobold, Redirecting engineered immune cells using G protein-coupled receptors in cancer therapy. Immuno-Oncology and Technology 29, 101582 (2026).

13. M. Serafini, B. Sacchetti, A. Pievani, D. Redaelli, C. Remoli, A. Biondi, M. Riminucci, P. Bianco, Establishment of bone marrow and hematopoietic niches in vivo by reversion of chondrocyte differentiation of human bone marrow stromal cells. Stem Cell Research 12, 659–672 (2014).

14. A. Abarrategi, S. A. Mian, D. Passaro, K. Rouault-Pierre, W. Grey, D. Bonnet, Modeling the human bone marrow niche in mice: From host bone marrow engraftment to bioengineering approaches. J Exp Med 215, 729–743 (2018).

15. A. Grigoryan, D. Zacharaki, A. Balhuizen, C. R. Côme, A. G. Garcia, D. Hidalgo Gil, A.-K. Frank, K. Aaltonen, A. Mañas, J. Esfandyari, P. Kjellman, E. Englund, C. Rodriguez, W. Sime, R. Massoumi, N. Kalantari, S. Prithiviraj, Y. Li, S. J. Dupard, H. Isaksson, C. D. Madsen, B. T. Porse, D. Bexell, P. E. Bourgine, Engineering human mini-bones for the standardized modeling of healthy hematopoiesis, leukemia, and solid tumor metastasis. Science Translational Medicine 14, eabm6391 (2022).

16. A. Reinisch, D. Thomas, M. R. Corces, X. Zhang, D. Gratzinger, W.-J. Hong, K. Schallmoser, D. Strunk, R. Majeti, A humanized bone marrow ossicle xenotransplantation model enables improved engraftment of healthy and leukemic human hematopoietic cells. Nat Med 22, 812–821 (2016).

17. M. Iafrate, G. O. Fruhwirth, How Non-invasive in vivo Cell Tracking Supports the Development and Translation of Cancer Immunotherapies. Front. Physiol. 11 (2020).

18. M. S. Skovgard, H. R. Hocine, J. K. Saini, M. Moroz, R. Y. Bellis, S. Banerjee, A. Morello, V. Ponomarev, J. Villena-Vargas, P. S. Adusumilli, Imaging CAR T-cell kinetics in solid tumors: Translational implications. Molecular Therapy - Oncolytics 22, 355–367 (2021).

19. A. G. T. Chavez, M. K. McKenna, K. Balasubramanian, L. Riffle, N. L. Patel, J. D. Kalen, B. S. Croix, A. M. Leen, P. Bajgain, A dual-luciferase bioluminescence system for the assessment of cellular therapies. Molecular Therapy Oncology 32 (2024).

20. Y. Su, J. R. Walker, Y. Park, T. P. Smith, L. X. Liu, M. P. Hall, L. Labanieh, R. Hurst, D. C. Wang, L. P. Encell, N. Kim, F. Zhang, M. A. Kay, K. M. Casey, R. G. Majzner, J. R. Cochran, C. L. Mackall, T. A. Kirkland, M. Z. Lin, Novel NanoLuc substrates enable bright two-population bioluminescence imaging in animals. Nat Methods 17, 852–860 (2020).

21. D. Zhang, E. Krimitza, K. Han, R. Su, D. J. Xu, J. R. Xu, Y. Gong, Y. Fan, Protocol to generate traceable CAR T cells for syngeneic mouse cancer models. STAR Protoc 5, 102898 (2024).

22. R. E. Sanchez-Pupo, J. J. Kelly, N. Shalaby, Y. Xia, F. M. Martinez-Santiesteban, J. Lau, I. E. Verriet, M. S. Fox, J. W. Hicks, J. D. Thiessen, J. A. Ronald, Imaging CRISPR-edited CAR-T cell therapies with optical and positron emission tomography reporters. Theranostics 16, 3227–3245 (2026).

23. M. C. Rotiroti, C. Buracchi, S. Arcangeli, S. Galimberti, M. G. Valsecchi, V. M. Perriello, T. Rasko, G. Alberti, C. F. Magnani, C. Cappuzzello, F. Lundberg, A. Pande, G. Dastoli, M. Introna, M. Serafini, E. Biagi, Z. Izsvák, A. Biondi, S. Tettamanti, Targeting CD33 in Chemoresistant AML Patient-Derived Xenografts by CAR-CIK Cells Modified with an Improved SB Transposon System. Molecular Therapy 28, 1974–1986 (2020).

24. M. P. Hall, J. Unch, B. F. Binkowski, M. P. Valley, B. L. Butler, M. G. Wood, P. Otto, K. Zimmerman, G. Vidugiris, T. Machleidt, M. B. Robers, H. A. Benink, C. T. Eggers, M. R. Slater, P. L. Meisenheimer, D. H. Klaubert, F. Fan, L. P. Encell, K. V. Wood, Engineered Luciferase Reporter from a Deep Sea Shrimp Utilizing a Novel Imidazopyrazinone Substrate. ACS Chem. Biol. 7, 1848–1857 (2012).

25. G. Gui, M. A. Bingham, J. R. Herzog, A. Wong-Rolle, L. W. Dillon, M. Goswami, E. Martin, J. Reeves, S. Kim, A. Bahrami, H. F. Degenhardt, G. Zaki, P. Divakar, E. C. Schrom, K. R. Calvo, C. S. Hourigan, K. D. Hansen, C. Zhao, Single-cell spatial transcriptomics reveals immunotherapy-driven bone marrow niche remodeling in AML. Science Advances 11, eadw4871 (2025).

26. D. O. Treaba, D. M. Bonal, A. Chorzalska, M. Castillo-Martin, A. Oakes, M. Pardo, M. Petersen, C. Schorl, K. Hopkins, D. Melcher, T. C. Zhao, O. Liang, E.-Y. So, J. Reagan, A. J. Olszewski, J. Butera, D. C. Anthony, P. Rintels, P. Quesenberry, P. M. Dubielecka, Transcriptomics of acute myeloid leukaemia core bone marrow biopsies reveals distinct therapy response-specific osteo-mesenchymal profiles. Br J Haematol 200, 740–754 (2023).

27. L. Chen, E. Pronk, C. van Dijk, Y. Bian, J. Feyen, T. van Tienhoven, M. Yildirim, P. Pisterzi, M. M. E. de Jong, A. Bastidas, R. M. Hoogenboezem, C. Wevers, E. M. Bindels, B. Löwenberg, T. Cupedo, M. A. Sanders, M. H. G. P. Raaijmakers, A Single-Cell Taxonomy Predicts Inflammatory Niche Remodeling to Drive Tissue Failure and Outcome in Human AML. Blood Cancer Discov 4, 394–417 (2023).

28. D. Andreu-Sanz, L. Gregor, E. Carlini, D. Scarcella, C. Marr, S. Kobold, Predictive value of preclinical models for CAR-T cell therapy clinical trials: a systematic review and meta-analysis. J Immunother Cancer 13, e011698 (2025).

29. B. Sacchetti, A. Funari, S. Michienzi, S. Di Cesare, S. Piersanti, I. Saggio, E. Tagliafico, S. Ferrari, P. G. Robey, M. Riminucci, P. Bianco, Self-Renewing Osteoprogenitors in Bone Marrow Sinusoids Can Organize a Hematopoietic Microenvironment. Cell 131, 324–336 (2007).

30. A. Pievani, S. Donsante, C. Tomasoni, A. Corsi, F. Dazzi, A. Biondi, M. Riminucci, M. Serafini, Acute myeloid leukemia shapes the bone marrow stromal niche *in vivo*. Haematologica 106, 865–870 (2021).

31. C. Sauter, D. Zacharaki, A. G. Garcia, D. H. Gil, Y. Li, E. J. Pørtner, A. Grigoryan, M. Gabriel, A. Baudet, S. G. Antón, A. C. Simonsen, J. Cammenga, V. Lazarevic, P. E. Bourgine, Engineering autologous ossicles for the personalized modeling of acute myeloid leukemia. bioRxiv [Preprint] (2026). 10.64898/2026.04.13.715606.

32. T. Qi, K. McGrath, R. Ranganathan, G. Dotti, Y. Cao, Cellular kinetics: A clinical and computational review of CAR-T cell pharmacology. Adv Drug Deliv Rev 188, 114421 (2022).

33. S. J. Dupard, A. Grigoryan, S. Farhat, D. L. Coutu, P. E. Bourgine, Development of Humanized Ossicles: Bridging the Hematopoietic Gap. Trends Mol Med 26, 552–569 (2020).

34. A. Pievani, B. Sacchetti, A. Corsi, B. Rambaldi, S. Donsante, V. Scagliotti, P. Vergani, C. Remoli, A. Biondi, P. G. Robey, M. Riminucci, M. Serafini, Human umbilical cord blood-borne fibroblasts contain marrow niche precursors that form a bone/marrow organoid in vivo. Development 144, 1035–1044 (2017).

35. M. Tufail, C.-H. Jiang, N. Li, Immune evasion in cancer: mechanisms and cutting-edge therapeutic approaches. Sig Transduct Target Ther 10, 227 (2025).

36. C. Ashmore-Harris, M. Iafrate, A. Saleem, G. O. Fruhwirth, Non-invasive Reporter Gene Imaging of Cell Therapies, including T Cells and Stem Cells. Mol Ther 28, 1392–1416 (2020).

37. M. Biondi, S. Tettamanti, S. Galimberti, B. Cerina, C. Tomasoni, R. Piazza, S. Donsante, S. Bido, V. M. Perriello, V. Broccoli, A. Doni, F. Dazzi, A. Mantovani, G. Dotti, A. Biondi, A. Pievani, M. Serafini, Selective homing of CAR-CIK cells to the bone marrow niche enhances control of the acute myeloid leukemia burden. Blood 141, 2587–2598 (2023).

