## Supplementary Figures and Table for "A humanized ossicle platform for real-time tracking and preclinical assessment of CAR-based immunotherapies in acute myeloid leukemia"

### Supplementary Materials

#### Supplementary Fig.1

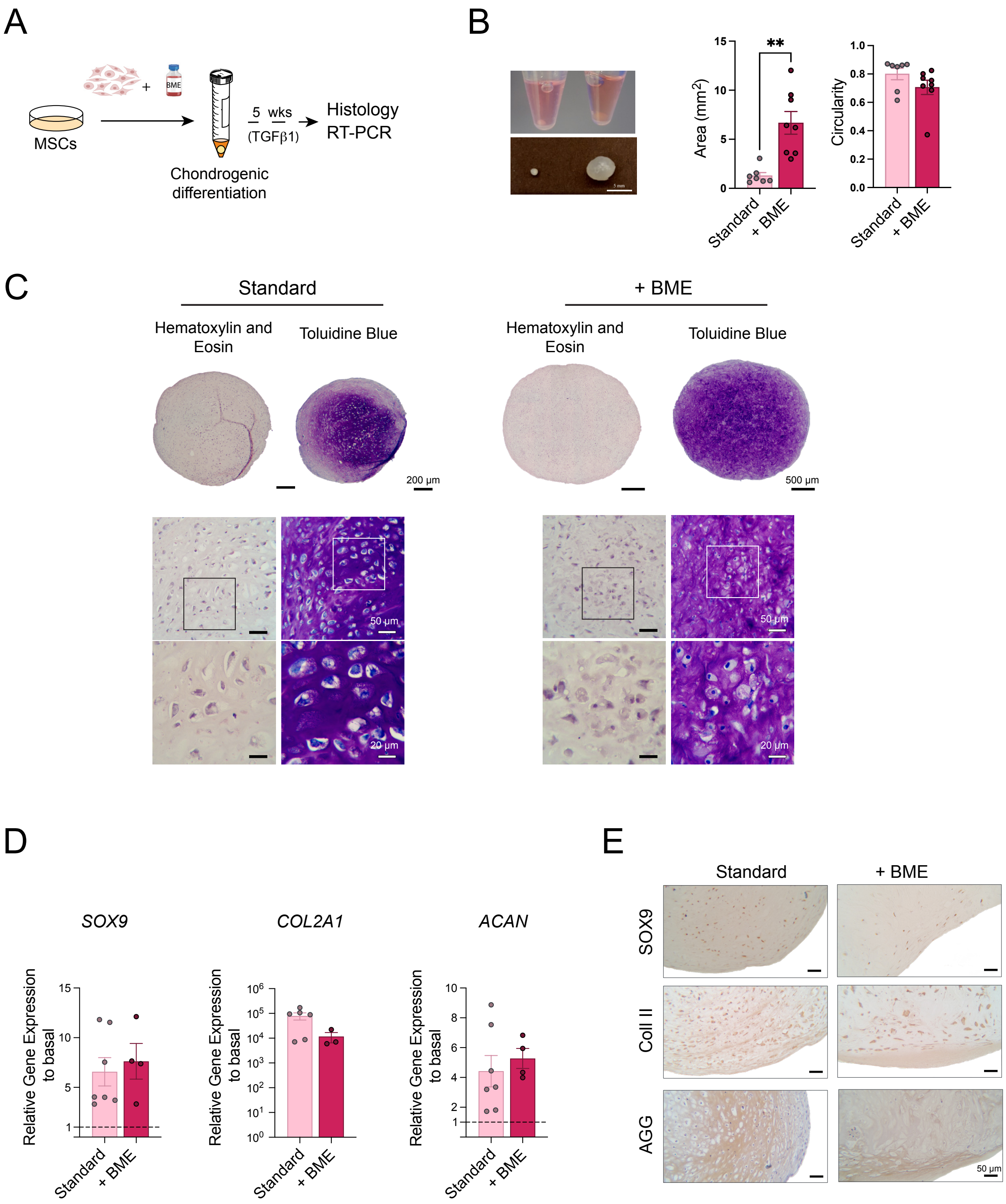


**Fig. S1. *In vitro* characterization of BME-cartilaginous pellets.**

**(A)** Schematic overview of BME-cartilaginous pellet generation from human BM mesenchymal stromal cells (MSCs). **(B)** Macroscopic view of standard (left) and BME-cartilaginous pellets (right).

Scale bar = 5mm. Quantification of area ($mm^2$) and circularity for standard (n=7) and BME-cartilaginous pellets (n=8). Data were derived from n=3 independent donors. Statistical values were determined by unpaired t test. (**C)** Hematoxylin and eosin and Toluidine Blue staining of paraffin sections from standard and BME-cartilaginous pellets. Scale bars = 200µm and 500µm for whole sections of standard and BME-pellets, respectively; 50µm and 20µm for zoomed areas. (**D)** Real-time PCR analysis of the chondrogenic markers *SOX9, COL2A1*, and *ACAN* in standard (n=7) and BME-cartilaginous pellets (n=4), relative to pre-differentiation MSC controls (n=4). Data were derived from n=3 independent donors. No significant differences were found by unpaired t test. **(E)** Representative immunohistochemical staining for SOX9, Collagen type II (Coll II) and Aggrecan (AGG) on paraffin sections of the pellets. Scale bar = 50µm.

**Supplementary Fig.2**

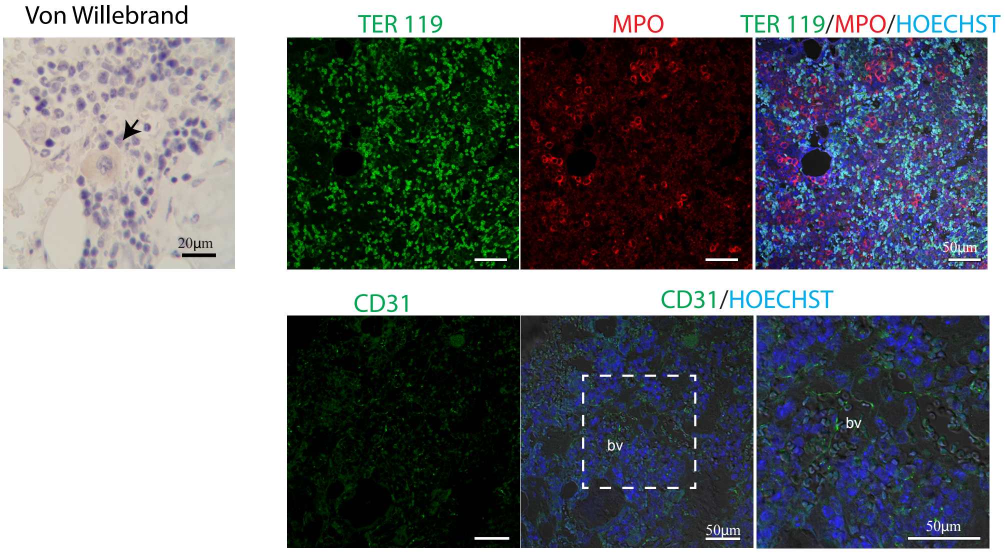


**Fig. S2. Characterization of murine components in BME-ossicles.**

Representative immunostaining for von Willebrand factor (megakaryocytes), double immunofluorescence for TER-119 (erythroid cells) and MPO (myeloid cells), and immunofluorescence for CD31 (endothelial cells) within the murine-derived hematopoietic compartment of BME-ossicles (MPO: myeloperoxidase). Nuclei were stained with Hoechst. Scale bar =20 and 50µm. bv: blood vessels.

**Supplementary Fig.3**

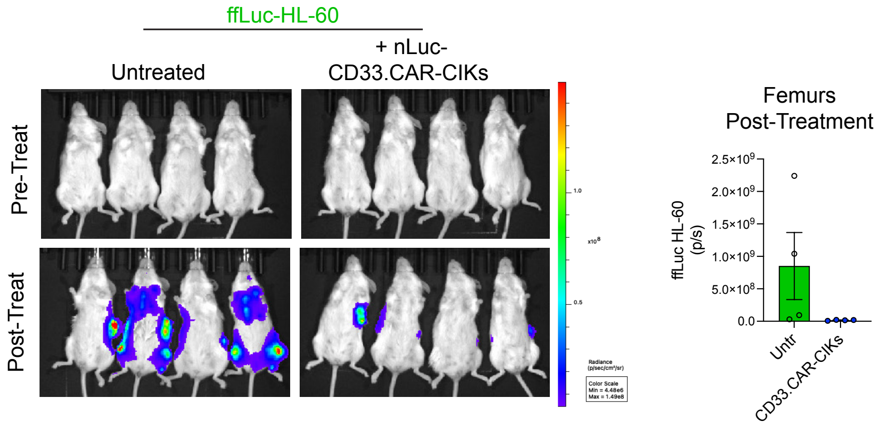


**Fig. S3.** ***In vivo*** **distribution of ffLuc-HL-60 cells before and after nLuc-CD33.CAR-CIK cells injection.**

Left: Representative BLI images showing ffLuc-HL-60 engraftment in femurs before treatment (top) and 14 days after nLuc-CD33.CAR-CIK cell infusion (bottom) in treated versus untreated groups. Right: Comparison of leukemia burden in mouse femurs between untreated and treated groups at 14 days post-treatment, quantified by BLI (total flux: photons/sec). n=4 mice per group.

**Supplementary Table 1: Primer sequences used for real-time PCR.**

| Gene: | | Sequence: 5'-3' |
|---|---|---|
| *SOX9* | Forward | GGCAAGCTCTGGAGACTTCTG |
| | Reverse | CCCGTTCTTCACCGACTTCC |
| *ACAN* | Forward | GATGATCTGGCACGAGAAGGG |
| | Reverse | CGTTTGTAGGTGGTGGCTGTG |
| *COL2A1* | Forward | AGATGACGGTCCCTCTGGTG |
| | Reverse | ATCCTCTCTCACCACGTTGC |
| *GAPDH* | Forward | GTCTCCTCTGACTTCAACAGCG |
| | Reverse | ACCACCCTGTTGCTGTAGCCAA |
